# Mitochondrial replacement in an iPSC model of Leber’s hereditary optic neuropathy

**DOI:** 10.1101/120659

**Authors:** Raymond C.B. Wong, Shiang Y. Lim, Sandy S.C. Hung, Stacey Jackson, Shahnaz Khan, Nicole J. Van Bergen, Elisabeth De Smit, Helena H. Liang, Lisa S Kearns, Linda Clarke, David A. Mackey, Alex W. Hewitt, Ian A. Trounce, Alice Pébay

**Author notes:** co-senior authors (Ian A. Trounce, Alice Pébay). Corresponding authors: Raymond C.B. Wong, Alice Pébay.

## Abstract

Cybrid technology was used to replace Leber hereditary optic neuropathy (LHON) causing mitochondrial DNA (mtDNA) mutations from patient-specific fibroblasts with wildtype mtDNA, and mutation-free induced pluripotent stem cells (iPSCs) were generated subsequently. Retinal ganglion cell (RGC) differentiation demonstrates increased cell death in LHON-RGCs and can be rescued in cybrid corrected RGCs.

## Introduction

Patient somatic cells can be reprogrammed into iPSCs, by ectopic expression of a combination of reprogramming factors (Takahashi et al., 2007), (Yu et al., 2007). These cells can subsequently be differentiated into cell types of interest, providing a platform for disease modeling, drug screening and gene therapy. In recent years, the field of iPSC disease modeling has focused on the use of isogenic control, *i. e*. by correcting the disease-causing mutation(s) to verify the disease phenotypes observed in the iPSC model. Gene editing technologies, such as TALEN or CRISPR/Cas9, emerged as tools to establish isogenic iPSC controls for nuclear gene mutations. However, the difficulty of genetically manipulating mtDNA represents an obstacle for generating iPSC isogenic controls to model mtDNA diseases. Mutations in mtDNA can be heteroplasmic (mutated mtDNA coexist with wild-type mtDNA) or homoplasmic (all mtDNA copies are mutated). iPSCs have been utilised to model mitochondrial diseases with mtDNA heteroplasmy (Xu et al., 2013). Interestingly, mutated mtDNA heteroplasmy has been reported to segregate during iPSC reprogramming, similar to the mtDNA genetic bottleneck phenomenon observed during early embryogenesis. In Mitochondrial Encephalomyopathy Lactic Acidosis and Stroke-like episodes (MELAS) (Folmes et al., 2013), Pearson Syndrome (Cherry et al., 2013) and diabetes mellitus (Fujikura et al., 2012), iPSC reprogramming yielded both mutant mtDNA-rich iPSC clones and mutation-free iPSC clones. However, this approach would not be feasible for iPSC modeling of homoplasmic mtDNA disease, such as LHON.

LHON is characterized by loss of the RGCs (Lopez Sanchez, Crowston, Mackey, & Trounce, 2016). In European populations, approximately 1 in 9,000 is a LHON carrier and it affects approximately 1 in 30,000 individuals, causing sudden visual loss predominantly (Yu-Wai-Man, Griffiths, & Chinnery, 2011). All LHON cases are caused by mtDNA mutations that encode for the mitochondrial Complex I subunits (Wallace et al., 1988), (Mackey et al., 1996). These homoplasmic mutations are shown to disrupt the activity of Complex I, leading to a decrease in bioenergetic production and an increased level of oxidative stress (Kirches, 2011). However, the precise mechanism for disease progression in LHON remains unknown. Here we report the use of iPSCs to model LHON, and demonstrate generation of isogenic iPSC controls by replacing LHON mtDNA using cybrid technology.

## Results

We previously reported on the generation and characterisation of LHON iPSCs (Hung et al., 2016). For this study, we utilised iPSCs from a healthy control (MRU11780), and a LHON patient (LHON Q1-4) with homoplasmic double mtDNA mutations m.4160T>C and m.14484T>C which affected the *MT-ND1* and *MT-ND6* genes respectively (Hung et al., 2016). This LHON patient exhibited “LHON plus” phenotype, with clinical features including optic nerve atrophy, juvenile encephalopathy and peripheral neuropathy (Mackey, 1994). To generate an isogenic control for iPSC modeling, we utilised the cybrid technique to replace the mutant mtDNA in LHON fibroblasts. LHON fibroblasts were pre-treated with rhodamine 6-G to disable the transmission of endogenous mtDNA, followed by fusion with donor mitochondria obtained from wild-type keratinocytes (Fig. 1A). After 27 days post-fusion, proliferating fibroblast colonies that are indicative of successful mitochondrial replacement were isolated and expanded (Fig. 1B). In contrast, no proliferating fibroblast colony was observed in control condition that did not receive donor keratinocyte mitochondria (Fig. 1B). Out of 12 clones screened, we identified 1 cybrid clone with the corrected mtDNA genotype at m.4160 and m.14484 (Fig. 1C). Microsatellite analysis of a panel of 12 polymorphic markers confirmed that the corrected cybrid clone originated from the parental LHON fibroblasts, whereas the donor keratinocytes possessed a different microsatellite profile (Fig. 1D, Supplementary File 1). We then generated iPSCs from the cybrid fibroblasts using the episomal method. Following reprogramming, three clones of cybrid iPSCs (CYB iPSC c1, CYB iPSC c2, CYB iPSC c3) were selected for this study. Importantly, all cybrid iPSC clones retained mtDNA correction with no detectable mutations at m.4160 and m.14484 (Fig. 1C, Supplementary File 2A). Further characterization demonstrated that the cybrid iPSCs expressed the pluripotent markers OCT-4 and TRA-1-60 (Fig. 1E, Supplementary File 2B). The derived cybrid iPSCs can also differentiated into cells of the three germ layers *in vitro* by embryoid body formation and *in vivo* by teratoma formation (Fig. 1F, Supplementary File 2B, C). Copy number variation analysis indicated no chromosomal abnormalities in the derived cybrid iPSC clones (Fig. 1G, Supplementary File 2D). Together, these results demonstrate the feasibility of using the cybrid technique to generate isogenic iPSC controls for mtDNA disease modeling.

**Figure 1.**
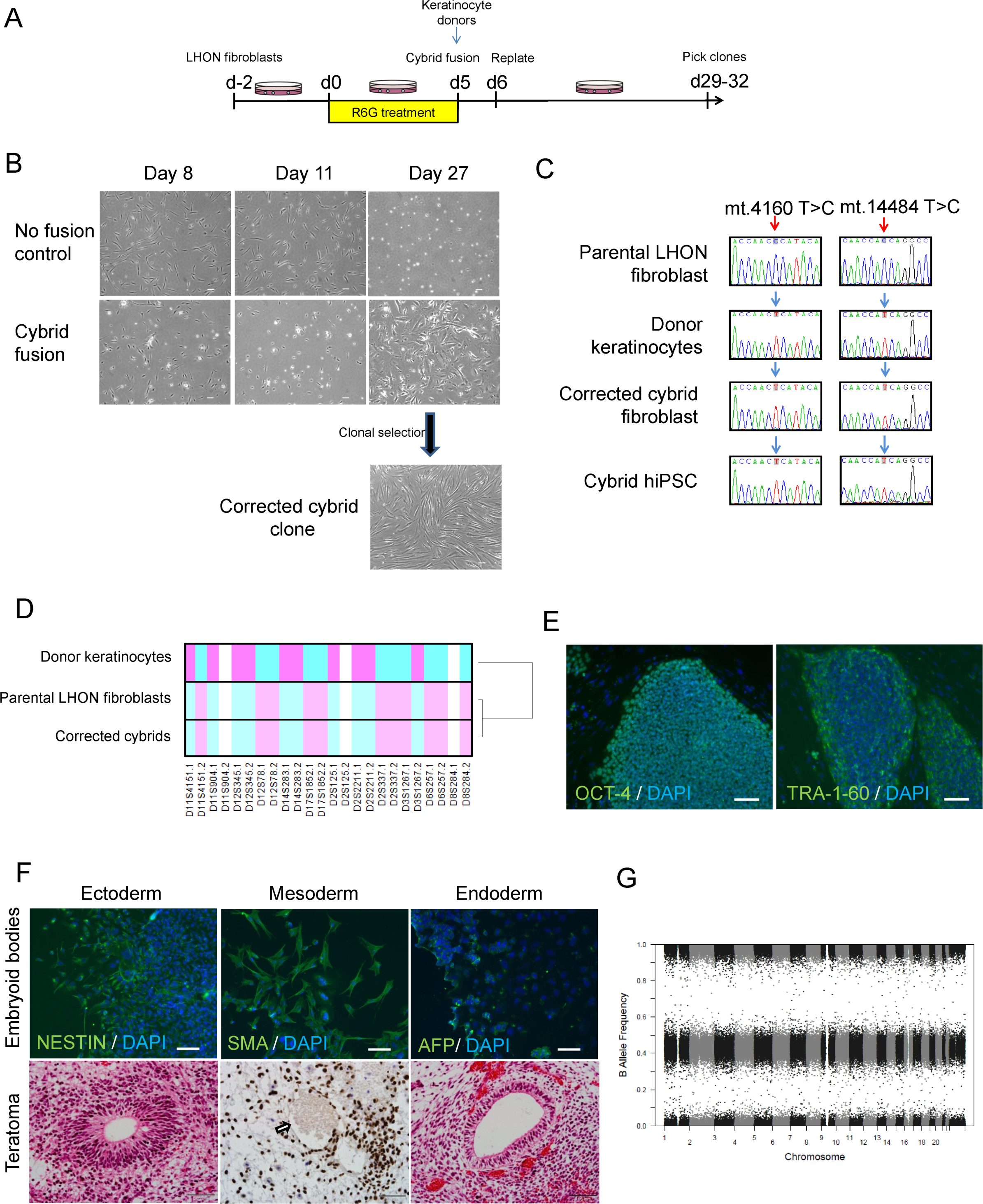
Using cybrid transfer to generate mutation-free LHON fibroblasts and iPSCs. **(A)**Diagram of cybrid generation. Fibroblasts were pre-treated with the mitochondrial toxin rhodamine 6-G (R6G) then fused with healthy donor mitochondria obtained from normal keratinocytes. On day 29-32, proliferating colonies were picked and expanded. **(B)** Representative images of control fibroblasts (no fusion) and fused fibroblasts that received donor mitochondria (cybrid fusion) at 8, 11, 27 days post R6G treatment. **(C)** Genotype confirming cybrid correction of mutation in fibroblasts and the corresponding iPSCs. Red arrows indicated LHON mutations at m.4160T>C and m.14484T>C, blue arrows indicate wild-type genotype. Note that genotype of parental LHON fibroblast (LHON Q1-4) was reported previously (Hung et al., 2016). **(D)** Microsatellite analysis confirming cybrid originated from LHON fibroblasts, but not donor keratinocytes. **(E-G)** Characterization of cybrid iPSCs (CYB iPSC c1). **(E)** Immunostaining showed expression of the pluripotency markers OCT-4 and TRA-1-60 in cybrid iPSCs. Scale bars: 100 µm. **(F)** Top panel: Differentiation of cybrid iPSCs by embryoid body formation contained cells positive for NESTIN (ectoderm), SMA (mesoderm) and AFP (endoderm) expression. Cells were counterstained with DAPI (blue). Scale bars: 100 µm. Bottom panel: Teratoma formation upon transplantation of cybrid iPSCs in nude rats, showing differentiation to endoderm, mesoderm (Ku80 staining, arrow indicate endothelial-lined blood vessel with lumen filled with red blood cells) and ectoderm. Scale bars: 50 µm. **(G)** Copy number variation analysis showing normal karyotype in cybrid iPSCs.

Finally, we assessed the effect of the LHON mtDNA mutations in iPSC-derived RGCs. Control, LHON and cybrid iPSCs were directed to differentiate into RGCs by a stepwise differentiation method that we recently published, which demonstrated an enriched population of functional RGCs (Gill et al., 2016). Three iPSC clones per patient were used and all clones were able to differentiate into RGCs with similar efficiency (Fig. 2A). Following RGC enrichment using MACS, TUNEL analysis revealed an increased level of apoptosis in LHON RGCs, from 13.6 ± 1.9% in control RGCs to 56.1 ± 6.8% in LHON RGCs (Fig. 2B). Importantly, this effect was reverted in the cybrid corrected RGCs, with the apoptosis level returned to control levels (Fig. 2B, 12.9 ± 4.4%), demonstrating that the increased susceptibility to cell death observed in LHON-RGCs is a direct consequence of LHON mtDNA mutations. Mitochondrial dysfunction in RGCs was then assessed using MitoSOX, which measures the levels of mitochondrial superoxide, as an indication of mitochondrial oxidative stress. Despite a trend of an increased superoxide levels in LHON RGCs compared to control RGCs, this difference in mitochondrial superoxide levels was not statistically significant, possibly due to high variations observed amongst the iPSC clones. Lower superoxide levels were also observed in corrected cybrid lines compared to LHON RGCs (Fig. 2C). Together these results suggest that the higher apoptosis observed in the LHON RGCs cannot be explained by elevated mitochondrial superoxide alone. Future studies to investigate other defects in LHON RGCs, such as ATP deficiency (Baracca et al., 2005) or mitochondrial biogenesis (Giordano et al., 2014), will help elucidate the mechanism underlying penetrance and disease progression of LHON.

**Figure 2.**
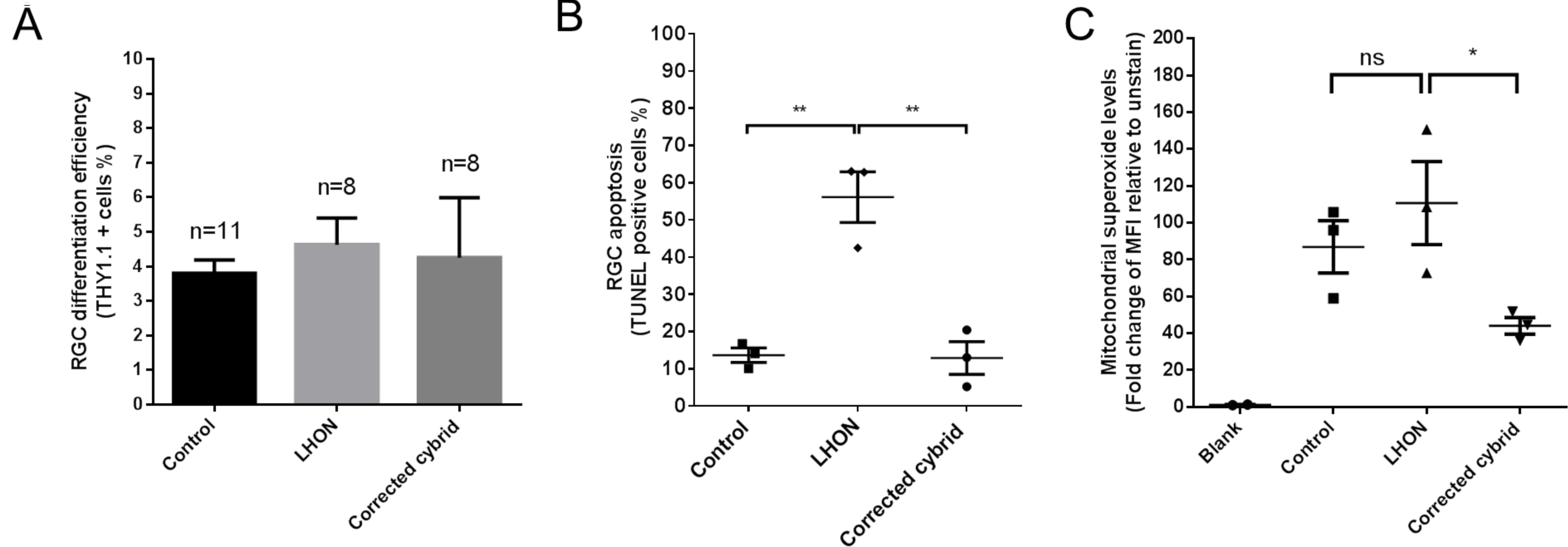
LHON disease phenotype in iPSC-derived RGCs can be reversed in corrected cybrid lines. **(A)** Efficiency in RGC differentiation as assessed by the % of THY1.1 positive cells obtained post MACS sorting for control, LHON and corrected cybrid lines. **(B)** TUNEL assay revealed increased susceptibility to apoptosis in LHON RGCs and reversal in corrected cybrid lines. Data are expressed as mean of each clone, n = 3 clones, error bars = mean ± SEM. **(C)** Quantification of mitochondrial superoxide levels using MitoSOX in control, LHON and corrected cybrid RGCs. Error bars = ± SEM, n = 3 clones. Statistical significance was established by one way ANOVA followed by Dunnett’s test for multiple comparisons, ** p<0.01, * p<0.05, ns: not significant.

## Discussion

The recent development of mitochondrial replacement therapy using pronuclear transfer offer exciting prospects to correct mtDNA mutations in the early human embryo (Hyslop et al., 2016). However, pronuclear transfer is technically challenging, especially in cells with small sizes, and requires specialised equipment. Here, we describe for the first time a cybrid approach to correct mtDNA mutations in an iPSC disease model. Compared to pronuclear transfer, the cybrid technique is easier to perform and can be adapted to generate an isogenic control for iPSC models of homoplasmic mtDNA diseases. Reduction of mutant mtDNA load, rather than complete replacement of mutant mtDNA with wild-type mtDNA, might be sufficient for phenotypic rescue in iPSC isogenic controls. In support of this, previous studies have shown that selective elimination of mutant mtDNA by mitochondria-targeted TALEN can reduce the mutant mtDNA loads and restore the mitochondrial dysfunction (Hashimoto et al., 2015). Very recently, zinc fingers were also used for the successful replacement of heteroplasmic mtDNA (Gammage et al., 2016). Our results show an increased susceptibility to apoptosis in LHON iPSC-derived RGCs. Importantly, apoptosis was returned to normal levels in the cybrid-corrected RGCs, hence demonstrating that the increased susceptibility to cell death observed in RGCs is a direct consequence of LHON mtDNA mutations.

In summary, our approach shows the advantage of using LHON patient-derived iPSCs and their isogenic cybrid control as a platform for the study of the fundamental mechanisms underlying LHON pathogenesis. Moreover, the cybrid technique provides a feasible strategy to correct mtDNA mutations in iPSC models, which could be applied to model other mtDNA diseases.

## Materials and Methods

### Ethics

All experimental work performed in this study was approved by the Human Research Ethics Committees of the Royal Victorian Eye and Ear Hospital (11/1031H) and the University of Melbourne (0605017) (McCaughey, Chen, et al., 2016; McCaughey, Liang, et al., 2016), and with the Animal Ethics Committee of St Vincent’s Hospital (002/14), in accordance with the requirements of the National Health & Medical Research Council of Australia and conformed with the Declarations of Helsinki.

### Cell culture

Cell lines used in this study consisted of control (MRU11780 (Hung et al., 2016)), LHON (LHON Q1-4 carrying m.14484 & m.4160 mtDNA mutations (Hung et al., 2016)) and cybrid-corrected lines. Fibroblasts were cultured in DMEM medium supplemented with 10% fetal calf serum, 1 × L-glutamine, 0.1 mM non-essential amino acids and 0.5× penicillin/streptomycin (all from Invitrogen). iPSC lines were cultured on mitotically inactivated mouse embryonic fibroblasts feeders in the presence of DMEM/F-12 medium supplemented with 1 × GlutaMAX, 20% knockout serum replacement, 10 ng/ml basic fibroblast growth factor, 0.1 mM non-essential amino acids, 100 µM β-mercaptoethanol and penicillin/streptomycin (all from Invitrogen) and passaged weekly as previously described (Hung et al., 2016).

### Cybrid generation

Cybrid transfer was performed as described (Trounce & Wallace, 1996; Williams, Murrell, Brammah, Minchenko, & Christodoulou, 1999). Briefly, human epidermal keratinocytes (System Bioscience) were used as donor by enucleation using cytochalasin B treatment and high speed centrifugation. LHON Q1-4 fibroblasts were pre-treated with 2.5µg/ml rhodamine 6-G for 5 days, before fusion with donor cell cytoplasts using polyethylene glycol. On the next day, fused cells were replated and allowed to culture for up to 32 days. Proliferating colonies were picked and expanded using cloning cylinders (Corning). Isolated fibroblast clones were genotyped for mtDNA mutations at mt.4160 and mt.14484 to screen for successful mitochondrial replacement.

### Microsatellite analysis

Microsatellite analysis was performed by the Australian Genomics Research Facility, using 12 polymorphic markers from the Applied Biosystems linkage mapping set. The markers utilised are D2S2211, D2S125, D2S337, D3S1267, D6S257, D8S284, D11S904, D11S4151, D12S78, D12S345, D14S283 and D17S1852. Hierarchical clustering and heatmap is generated using R.

### iPSC generation and characterisation

Reprogramming of cybrid-corrected fibroblasts was performed as described in (Piao, Hung, Lim, Wong, & Ko, 2014). Episomal vectors expressing OCT4, SOX2, KLF4, L-MYC, LIN28 and shRNA against p53 were gifts from Shinya Yamanaka (Addgene #27077, 27078, 27080). *In vitro* differentiation of iPSCs was performed by embryoid bodies and characterised as described in (Hung et al., 2016). *In vivo* teratoma assay was performed by transplanting iPSCs into a vascularized tissue engineering chamber in immuno-deficient rats, as described in (Piao et al., 2014). Histological analysis was performed on the teratoma samples after 4 weeks. Copy number variation (CNV) analysis of original fibroblasts and iPSCs was performed using Illumina HumanCore Beadchip arrays. CNV analyses were performed using PennCNV with default parameter settings (Wang et al., 2007). Chromosomal aberrations were deemed to involve at least 10 contiguous single nucleotide polymorphisms (SNPs) or a genomic region spanning at least 1MB (Kilpinen et al., 2016; Wang et al., 2007).

### RGC differentiation

iPSCs were directed for retinal differentiation and enriched for RGCs using MACS THY1.1 microbeads (Miltenyi Biotech) as described in our previous study (Gill et al., 2016). On day 30, RGC differentiation efficiency is determined as described previously (Gill et al., 2016).

### Immunochemistry

Standard immunochemistry procedure was performed using mouse anti-OCT3/4 (#SC-5279, Santa Cruz Biotechnology), mouse anti-TRA-1-60 (#MAB4360, Millipore), mouse anti-NESTIN (#AB22035, Abcam), mouse anti-alpha-fetoprotein (AFP, #ST1673, Millipore), mouse anti-smooth muscle actin (SMA, #MAB1420, R&D systems). Cells were then immunostained with the appropriate conjugated secondary antibodies (Alexa Fluor 488, Molecular probes-Invitrogen). Nuclei were counter-stained with DAPI (Invitrogen). Immunohistochemistry for human cells was performed using rabbit anti-Ku80 (#AB80592, Abcam) followed by biotinylated goat anti-rabbit secondary antibody (Vector Laboratories) and avidin-biotinylated-peroxidase complex (Vector Laboratories). Peroxidase activity was visualised with diaminobenzidine chromogen (Dako) and sections were counterstained with hematoxylin. Specificity of the staining was verified by the absence of staining in negative controls consisting of the appropriate negative control immunoglobulin fraction.

### Mitochondrial superoxide measurement

Quantification of mitochondrial superoxide was performed on day 30 iPSC-derived RGCs using MitoSOX (Invitrogen) according to the manufacturer’s instructions. Briefly, trypsinized cells were stained with Mitosox for 10 minutes at 37°C. Subsequently, samples were processed by flow cytometry using the MACSquant (Miltenyi). The median fluorescence intensity was determined using the MACSQuantify software and normalized to unstained control.

### Apoptosis assay

TUNEL assay was performed on day 37-41 iPSC-derived RGCs using the *In situ* Cell death detection kit (Roche) following manufacturer’s instructions. Floating apoptotic bodies were collected and the enriched RGCs were harvested by trypsinization. Quantitation of TUNEL positive cells was measured by flow cytometry using the MACSquant (Miltenyi).

### Statistical analysis

All statistical analyses and graphical data were generated using Graphpad Prism software (v5.04, www.graphpad.com) or in the R statistical environment (v3.1.2, https://cran.r-project.org/). Statistical methods utilised were One-way ANOVA followed Dunnett’s multiple comparisons test for multiple comparison. Statistical significance was established as * p<0.05 and **p<0.01.

## Abbreviations

Cybrid: cyb
induced pluripotent stem cells: iPSCs
Leber hereditary optic neuropathy: LHON
Mitochondrial Encephalomyopathy Lactic Acidosis and Stroke-like episodes: MELAS
mitochondrial: mt
retinal ganglion cells: RGCs

## Author Contributions

RCBW: concept and design, financial support, collection and/or assembly of data, data analysis and interpretation, manuscript writing, final approval of manuscript. SJ, SK, SYL, SSCH, NJvB, MD, EDS, HHL, LK, LC, DM: collection and/or assembly of data, data analysis and interpretation, final approval of manuscript. AWH, IAT: concept and design, financial support, data analysis and interpretation, final approval of manuscript. AP: concept and design, financial support, data analysis and interpretation, manuscript writing, final approval of manuscript.

## Conflicts of Interest

The authors declare no conflict of interest.

## Funding

This work was supported by grants from the National Health and Medical Research Council (NHMRC, RCBW, 1084256), the Australian Mitochondrial Disease Foundation (RCBW, AWH, IAT, AP), the Brockhoff foundation (IAT, AP) the University of Melbourne (RCBW, AP), and the Ophthalmic Research Institute of Australia (RCBW, NJVB, AP). AWH is supported by a NHMRC Practitioner Fellowship (APP1103329), AP by an Australian Research Council Future Fellowship (FT140100047), and RCBW by a Peggy and Leslie Cranbourne Foundation Fellowship as well as a Medical Advances Without Animals Trust Fellowship. The Centre for Eye Research Australia receives operational infrastructure support from the Victorian Government.

---

**Supplementary File 1:**
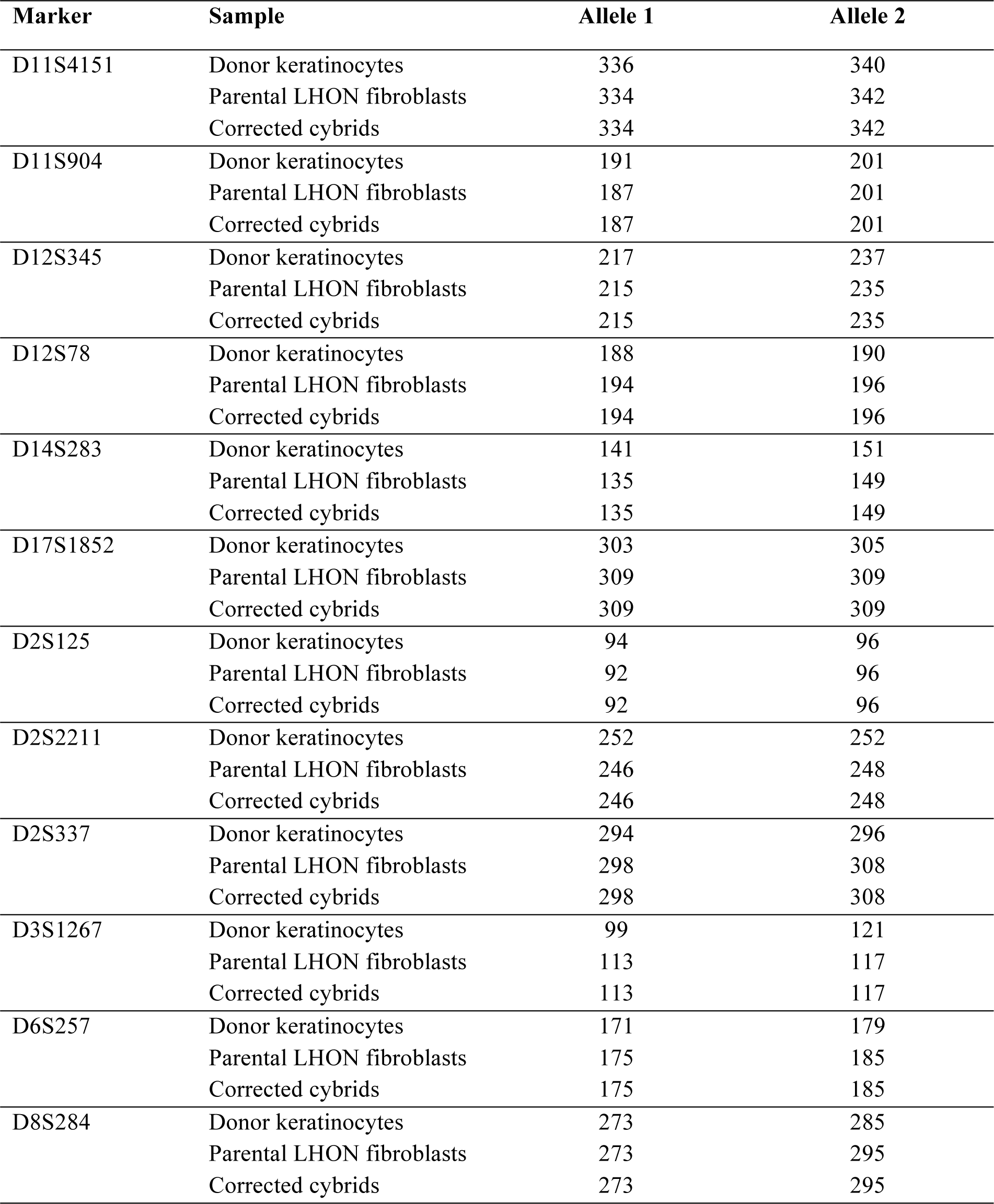
Profiles of 12 microsatellite markers for donor keratinocytes, parental LHON fibroblasts (LHON Q1-4) and corresponding corrected cybrids.

**Supplementary File 2A:**
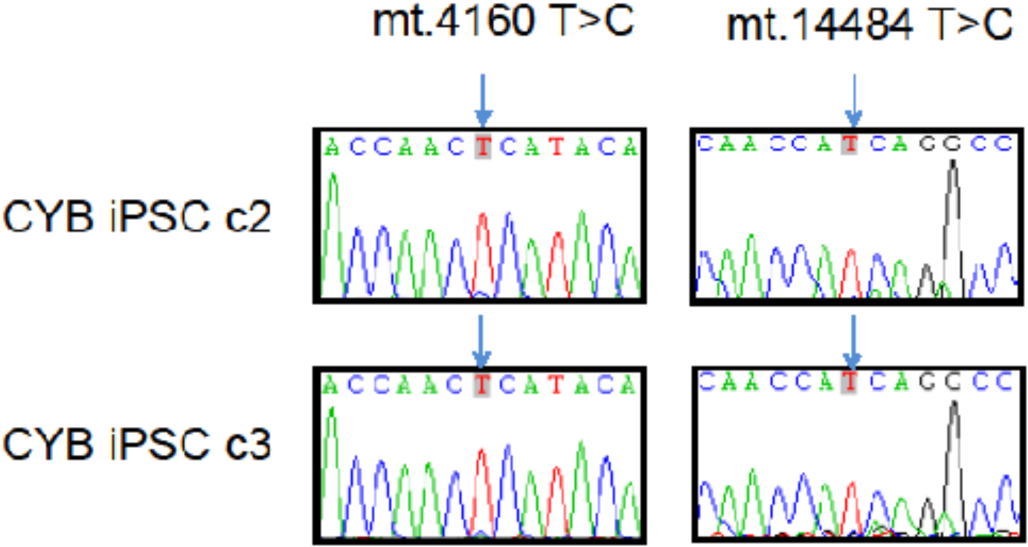
genotyping confirming two cybrid clones (CYB iPSC c2, CYB iPSC c3) with corrected mtDNA. Blue arrows indicate lack ok LHON mutations at m.4160T>C and m.14484T>C.

**Supplementary File 2B.**
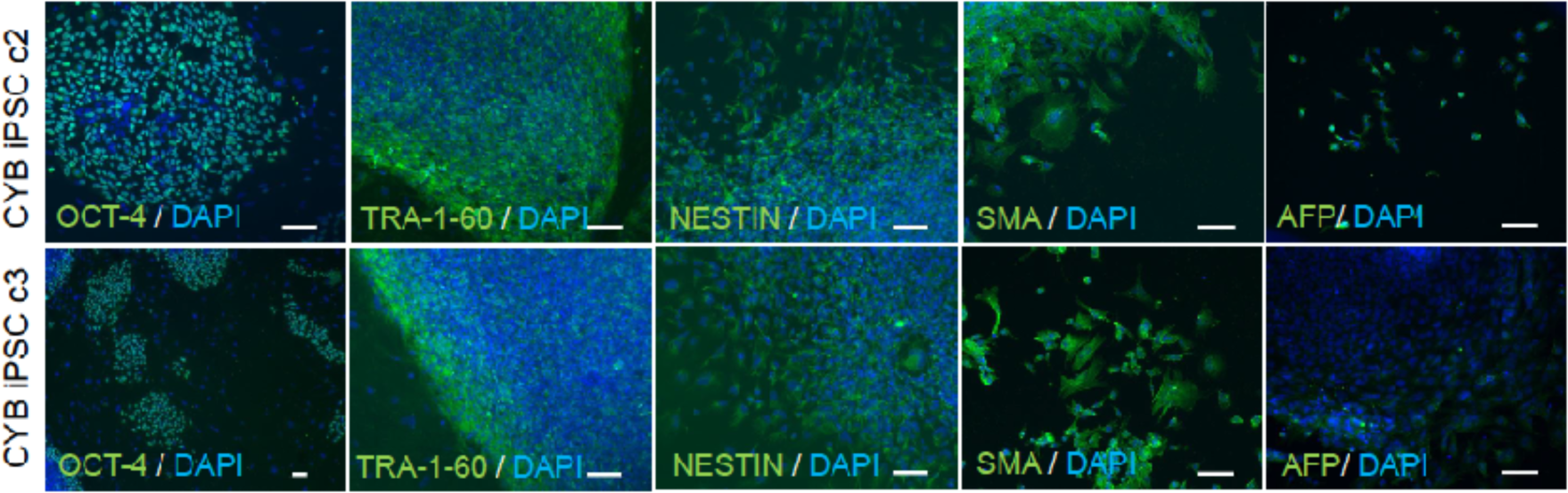
Characterization of two cybrid iPSCs (CYB iPSC c2, CYB iPSC c3). Immunostaining showed expression of the pluripotency markers OCT-4 and TRA-1-60 in cybrid iPSCs. Differentiation of cybrid iPSCs by embryoid body formation contained cells positive for NESTIN (ectoderm), SMA (mesoderm) and AFP (endoderm) expression. Cells were counterstained with DAPI (blue). Scale bars = 100 µm.

**Supplementary File 2C.**
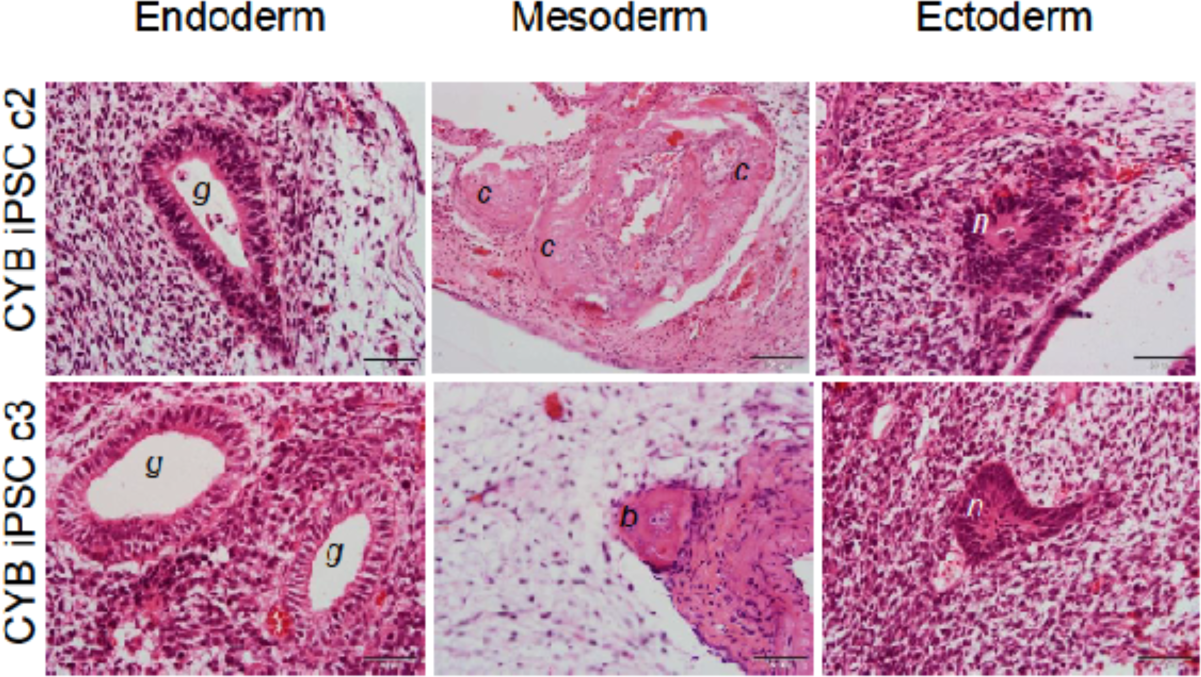
Teratoma formation upon transplantation of cybrid iPSCs (CYB iPSC c2, CYB iPSC c3) in nude rats, showing differentiation to endoderm, mesoderm and ectoderm. G: gut-like epithelium; c: cartilaginous structure; b: bone-like structure; n: neural rosette. Scale bars: 50 µm.

**Supplementary File 2D.**
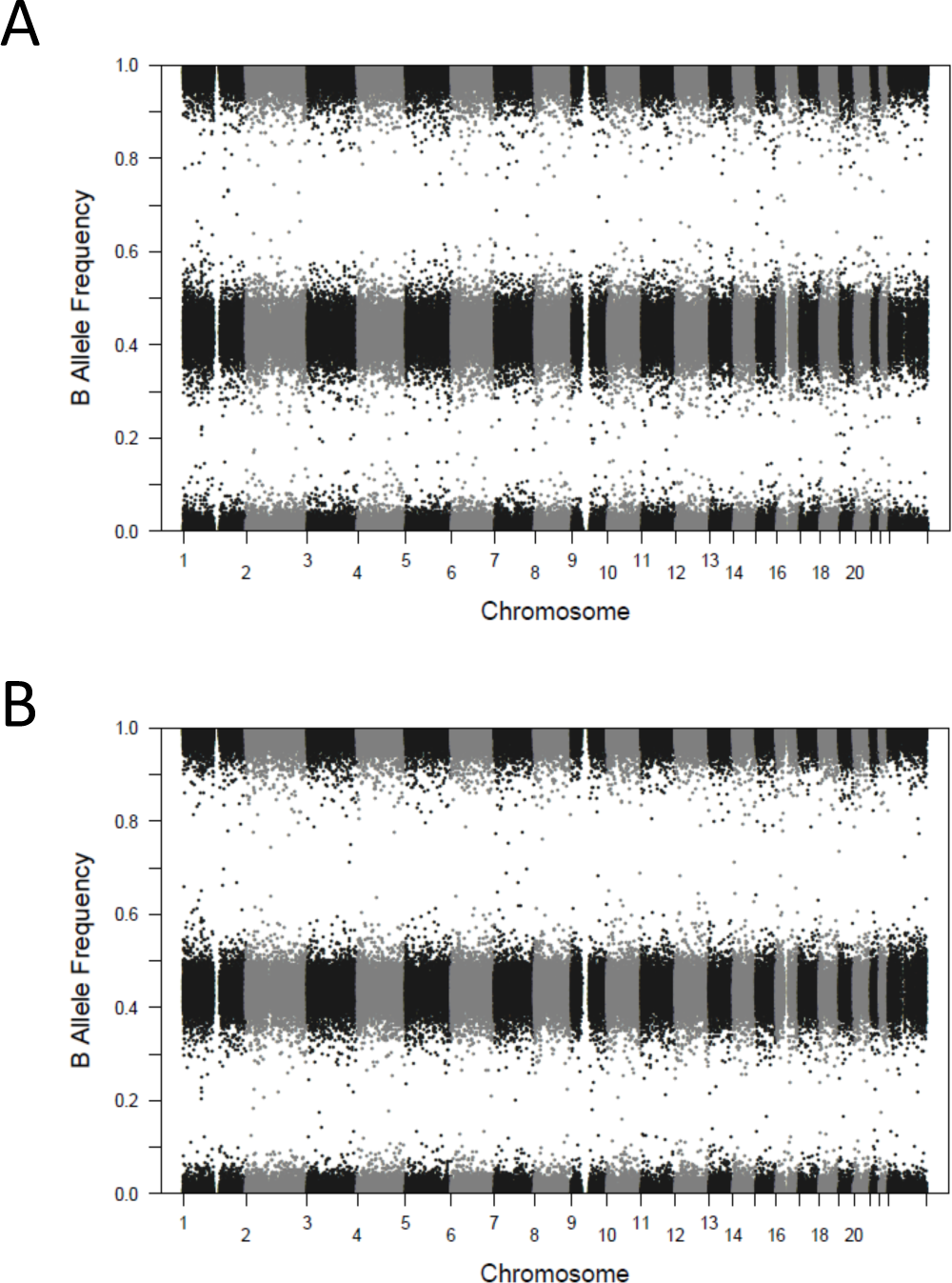
Copy number variation analysis showing normal karyotype in two cybrid iPSC clones, **(A)** CYB iPSC c2; **(B)** CYB iPSC c3.

